# A monoclonal antibody against staphylococcal enterotoxin B superantigen inhibits SARS-CoV-2 entry *in vitro*

**DOI:** 10.1101/2020.11.24.395079

**Authors:** Mary Hongying Cheng, Rebecca A. Porritt, Magali Noval Rivas, James M Krieger, Asli Beyza Ozdemir, Gustavo Garcia, Vaithilingaraja Arumugaswami, Bettina C. Fries, Moshe Arditi, Ivet Bahar

## Abstract

We recently discovered a superantigen-like motif, similar to Staphylococcal enterotoxin B (SEB), near the S1/S2 cleavage site of SARS-CoV-2 Spike protein, which might explain the multisystem-inflammatory syndrome (MIS-C) observed in children and cytokine storm in severe COVID-19 patients. We show here that an anti-SEB monoclonal antibody (mAb), 6D3, can bind this viral motif, and in particular its PRRA insert, to inhibit infection by blocking the access of host cell proteases, TMPRSS2 or furin, to the cleavage site. The high affinity of 6D3 for the furin-cleavage site originates from a poly-acidic segment at its heavy chain CDR2, a feature shared with SARS-CoV-2-neutralizing mAb 4A8. The affinity of 6D3 and 4A8 for this site points to their potential utility as therapeutics for treating COVID-19, MIS-C, or common cold caused by human coronaviruses (HCoVs) that possess a furin-like cleavage site.

---

Severe acute respiratory syndrome coronavirus 2 (SARS-CoV-2) can cause severe interstitial pneumonia with hyperinflammation^1,2^, as well as many extrapulmonary manifestations^3^. Furthermore, a novel multisystem inflammatory syndrome (MIS), reported in both children (MIS-C) and adults (MIS-A), has been observed in patients that either tested positive for, or had epidemiological links to, COVID-19^4-7^. MIS-C manifests as persistent fever and hyperinflammation with multi-organ system involvement^4-7^. The clinical similarity between MIS-C or severe COVID-19 and the toxic shock syndrome (TSS) caused by bacterial superantigens (SAg) led to the hypothesis that SARS-CoV-2 might possess a SAg-like motif that acts acutely or via an autoimmune-like mechanism to trigger hyperinflammation^8,9^. Comparison with bacterial toxins revealed a motif in the SARS-CoV-2 spike (S) protein whose sequence and structure are highly similar to a segment of a bacterial SAg, staphylococcal enterotoxin B (SEB). T cell receptor (TCR) skewing observed in severe COVID-19 patients further supported the SAg-like character of the S protein^8^.

The position of this putative SAg-like motif on the SARS-CoV-2 S structure is worthy of attention. SARS-CoV-2 S is a homotrimer, belonging to the family of human coronaviruses (HCoVs). This family includes SARS-CoV and Middle East Respiratory Syndrome (MERS), as well as the common cold HCoVs, NL63, 229E, OC43 and HKU1, that cause mild to moderate upper respiratory diseases^10-12^. Each HCoV-2 protomer is composed of two subunits, S1 and S2, playing different roles in viral infection. S1 contains the receptor-binding domain that binds to the receptor (angiotensin converting enzyme 2 (ACE2) in SARS-CoV-2, SARS-CoV, and HCoV-NL63) on the host cell^13-19^; S2 contains the fusion peptide required for viral entry^10-12^. The SAg-like motif lies at the C-terminus of S1 (residues E661-R685)^8^, at the boundary with S2. Membrane fusion requires two successive cleavages by host cell proteases, one at the S1/S2 interface (peptide bond R685-S686), and the other at S2’ (R815-S816)^10,13-18^. Thus, the SAg-like region overlaps with the S1/S2 cleavage site on the surface of the S glycoprotein (**Fig 1a-b**).

**Figure 1:**
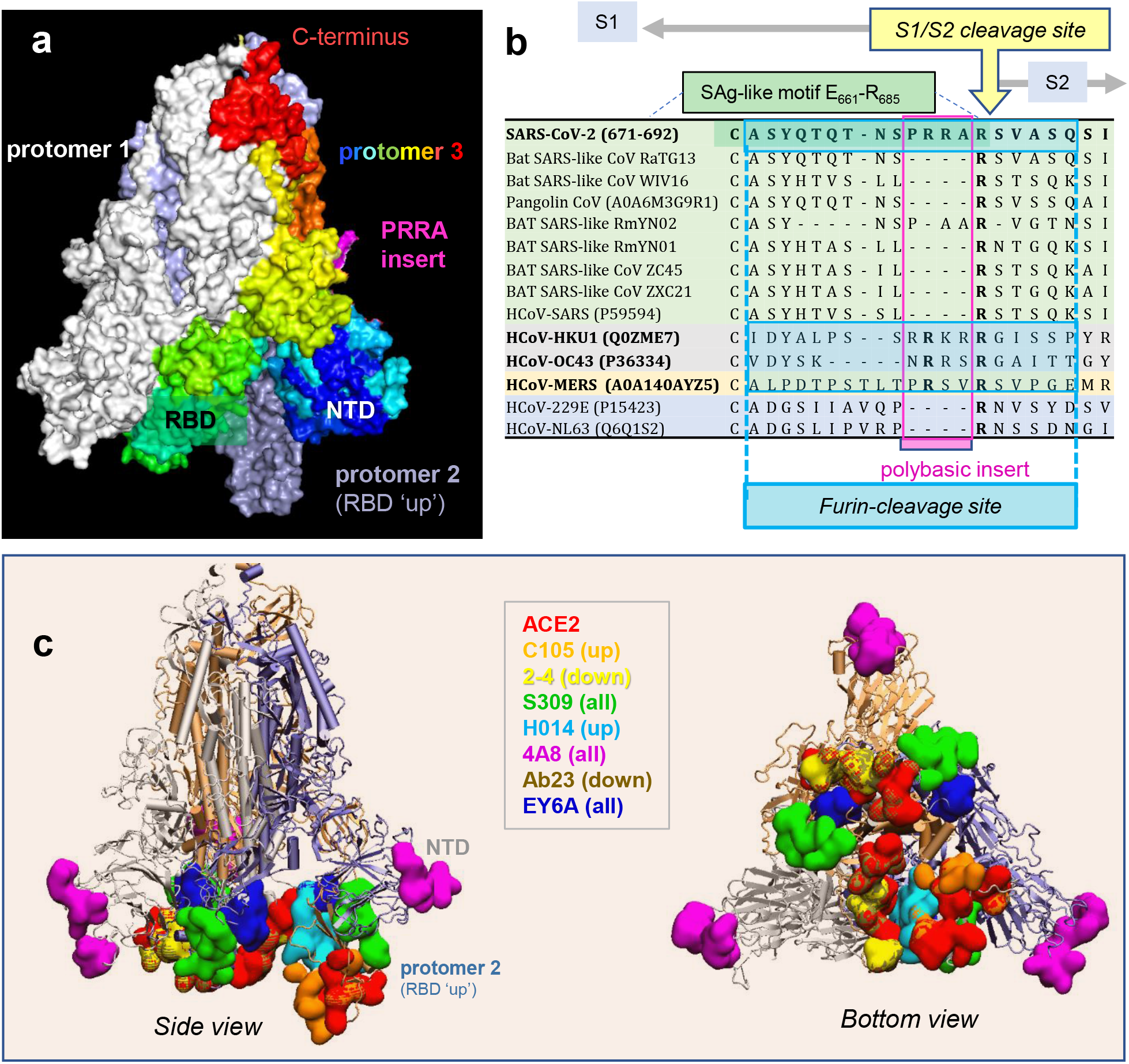
SARS-CoV-2 Spike (S) glycoprotein structure, sequence alignment against other CoVs, and interaction sites observed in cryo-EM studies with neutralizing antibodies. (**a**) SARS-CoV-2 S trimer in the pre-fusion state. Protomers 1 and 2 are in *white* and *light blue*, respectively; and protomer 3, in *spectral colors* from *blue* (N-terminal domain, NTD; residue 1-305) to *red* (C-terminus), except for the _681_PRR_A684_ insert in *magenta*. The insert is modeled using SWISS-MODEL^20^. Each protomer’s RBD (residues 331-524) can assume up or down conformations as indicated, linked to the respective receptor-bound and -unbound states. (**b**) Sequence alignment of SARS-CoV-2 near the S1/S2 cleavage site against multiple bat and pangolin SARS-related strains, and other HCoVs, adjusted following previous studies^10,21^. Viruses belonging to the same lineage are shown by the same *color shade*; and HCoVs that encode furin-like cleavage sites are highlighted in *bold fonts*. Note that the polybasic insert PRRA of SARS-CoV-2 S is not found in closely related SARS-like CoVs but exists in MERS and HCoVs HKU1 and OC43. The furin-like cleavage site is indicated by the *blue-shaded box*. (**c-d**) Side (*left*) and bottom (*right*) views of receptor (ACE2)- and antibody-binding sites observed in cryo-EM structures resolved for the S protein complexed with the ACE2 and/or various Abs. The S trimer is shown in *cartoons* with the light *blue* protomer in the RBD-up conformation, and *gray* and *light orange* protomers in the RBD-down conformation. Binding sites for ACE2 and antibodies C105^22^, 2-4^23^, S309^24^, H014^25^, 4A8^26^, Ab23^27^, and EY6A^28^ are shown in space-filling surfaces in different colors (see the code in the inset). See **Table 1** for additional information.

Another even more interesting feature near the S1/S2 cleavage site is the existence of a unique insertion, _681_PRRA_684_ (shortly PRRA) also taking part in the SAg-like motif and immediately neighboring the cleavage site R685-S686 (**Fig 1a**). SARS-CoV-2 is the only βCoV that has such an insertion, despite its high sequence similarity with other members of this genus (e.g. > 80% with SARS-CoV) (**Fig 1b**). Interestingly, MERS and common cold HCoVs HKU1 and OC43 S proteins also have a similar insertion at that position, despite their low (30-40%) overall sequence identity (**Fig 1b**). The PRRA insert is highly flexible, and together with the adjacent arginine, _681_PRRAR_685_ forms a highly reactive site. The insert is known to play a role in recognizing and binding the host cell proteases transmembrane protease serine 2 (TMPRSS2) and furin, whose cleavage activity is essential to S protein priming^14-17^. More recent studies further showed its role in facilitating SARS-CoV-2 entry and potentiating infectivity upon binding to the host cell co-receptor neuropilin-1^29,30^; and our simulations also pointed to its propensity to bind host cell TCRs^8^.

We hypothesized that such a polybasic site, _681_PRRAR_685_, implicated in key interactions with host cell proteins, could serve as a target for SARS-CoV-2 S-neutralizing antibodies (Abs). Most, if not all, SARS-CoV-2 S Abs currently under investigation are designed/developed to target the RBD (and some, the N-terminal domain, NTD)^24,26,27,31-34^. **Fig. 1c** illustrates the S-protein epitopes (*colored surfaces*) that have been observed by cryo-EM to bind mAbs and ACE2 molecules. These studies show that the Abs used as immunogens exhibit distinctive binding poses/sites favored by up (open) or down (closed) states of the RBDs (see **Table 1**). However, it is important to note that these cryo-EM studies were conducted in the presence of amino acid substitutions at the polybasic segment _682_RRAR_685_. The ability, if any, of the wild type (wt) S protein to bind an Ab near the PRRA insert or the S1/S2 cleavage may thus have eluded these experiments.

**Table 1:**
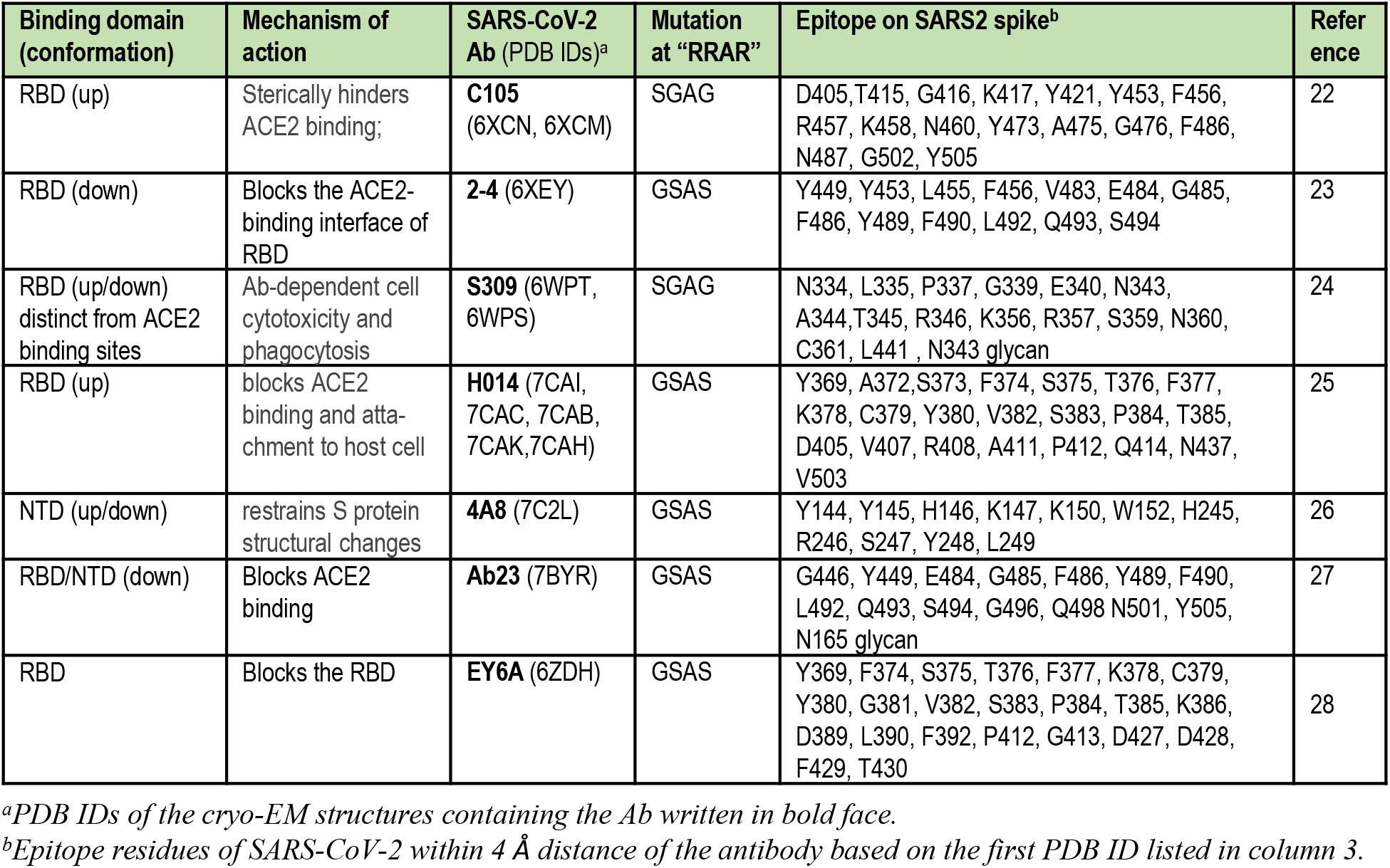
Antibody-bound complexes resolved by cryo-EM for SARS-CoV-2 Spike mutants.

We focus here on this polybasic site unique to SARS-CoV-2 among SARS-family βCoVs, as a target for mAb binding. In view of the recently detected sequence- and structure-similarity between the PRRA-insert-enclosing SAg-like motif and the bacterial enterotoxin SEB, we hypothesize that previously generated anti-SEB monoclonal Abs (mAbs) may potentially bind the viral SAg-like motif, and in particular the segment _682_RRAR_685_, and may thus block access to the S1/S2 cleavage site. We therefore examined *in silico* the possible interactions of known anti-SEB mAbs^35^ with SARS-CoV-2 S. Our study revealed the high affinity of SEB-specific mAb 6D3 for binding to the S1/S2 site. We also generated structural models for the interaction of the S-protein with TMPRSS2 and furin to demonstrate that the 6D3 binding site overlaps with those of these proteases, suggesting that 6D3 might interfere with viral entry. Experiments conducted with live viruses showed that 6D3 indeed inhibited viral entry to the host cell. Given that this site does not overlap with those observed to bind Abs (**Fig. 1c**), our study suggests that 6D3 might be used in combination with other neutralizing Abs that target the RBD or other non-overlapping sites to increase the efficacy of mAbs in inhibiting SARS-CoV-2 cellular entry.

Furthermore, we performed an *in silico* screening of SARS-CoV-2 neutralizing mAbs (**Table 1**), to probe their ability to bind near the PRRA insert, or the furin-like cleavage site. The latter typically contains eight central residues including the polybasic segment (here _680_SPRRAR↑SV_687_), flanked by solvent-accessible residues on both sides^36^. Our analysis revealed that 4A8^26^ may also bind the same site and as a result obstruct the S1/S2 site. Therefore, 4A8 and 6D3 may potentially serve as scaffold for designing wide-spectrum Abs for reducing the infectivity of SARS-CoV-2 and even other HCoVs that harbor a furin-like cleavage site (see **Fig. 1b**).

## Results

### TMPRSS2 and/or furin-bound to the S1/S2 site in close association with the PRRA insert

Host cell proteases cleave SARS-CoV-2 S at the S1/S2 site and induce a conformational change required for S protein priming for cell fusion and viral entry^14-18^. To assess whether Abs that might target the PRRA site would also hinder the access of proteases to the S1/S2 site, we first examined *in silico* the interaction between SARS-CoV-2 S protein and the proteases TMPRSS2 and furin. The resulting structural models are presented in the respective panels **a** and **b** of **Fig. 2**, and more details are reported in the Extended Data **Figs 1** and **2**. We used the available structural data^14,15,37^ for the three proteins, and the protein-protein docking software ClusPro^38^ and protocols outlined in the Supplemental Methods. An ensemble of structural models were generated for each complex, and those conformers satisfying the criteria for potential cleavage at the S1/S2 site, mainly close-positioning (within 3-7 Å atom-atom distance) of catalytic residues near the S1/S2 site, were selected for further refinement and energetic evaluation using PRODIGY^39^.

**Figure 2:**
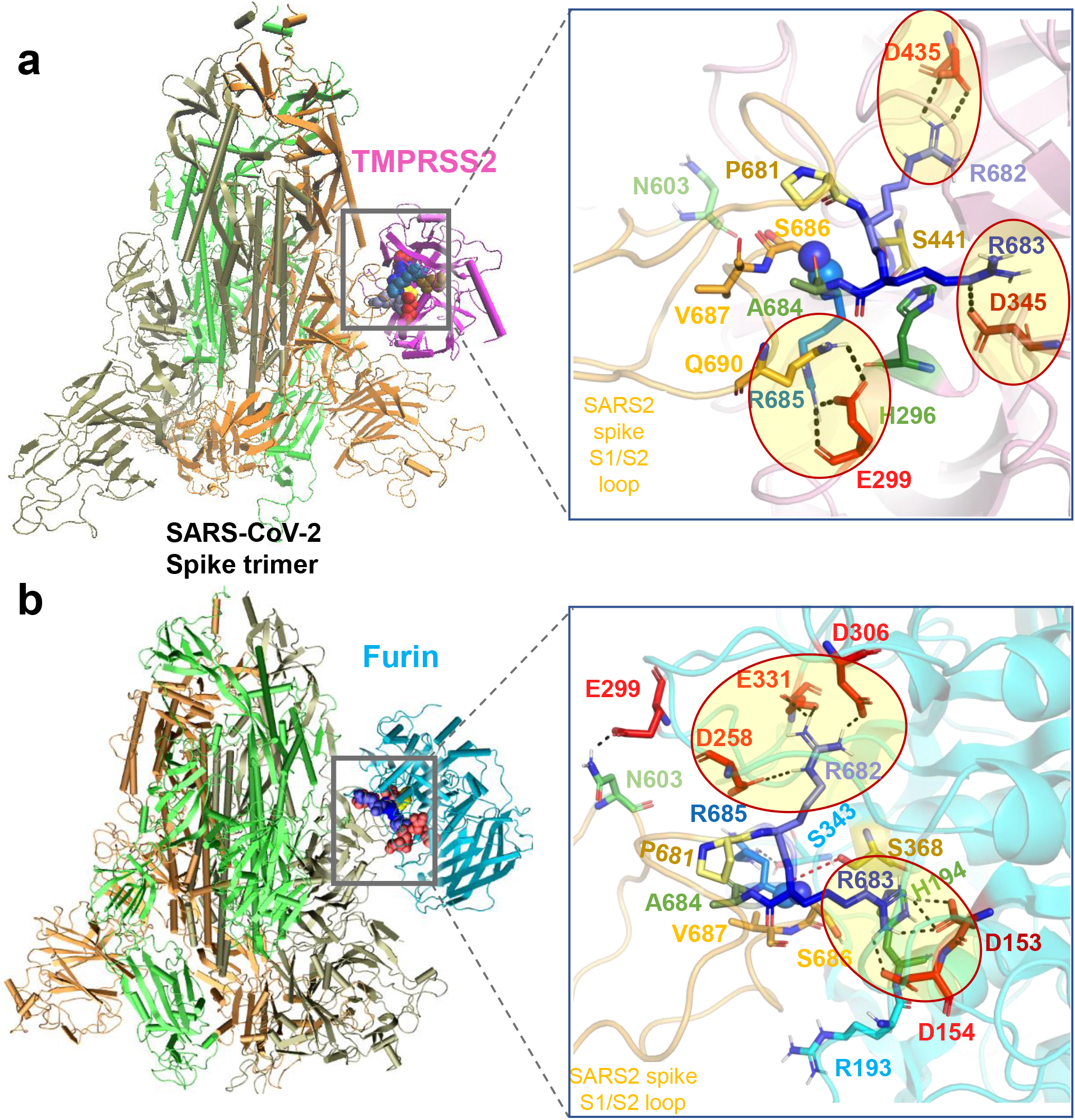
Binding poses of human proteases TMPRSS2 and furin to SARS-CoV-2 S protein. (**a-b**) Structural models for the SARS-CoV-2 S protein complexed with TMPRSS2 (**a**), and furin (**b**), obtained from docking simulations followed by refinements. An overview (*left*) and a zoomed in view (*right*) are shown in each case. The arginines in the S1/S2 loop P_681_RRARS_686_ are shown in different shades of *blue,* and their interaction partner (acidic residues) in the proteases are shown in *red*. *Spheres (right panels)* highlight the peptide bond that would be cleaved (between R685 and S686). TMPRSS2 catalytic triad residues are in *yellow* for S441, *green* for H296, and *dark* red for D345. Their counterparts in furin are S368, H194 and D153. Note the short distance between the carbonyl carbon from R685 and the hydroxyl oxygen of the catalytic serine S441 of TMPRSS2 (3.5 Å) or S368 of furin (3.1 Å). *Black dashed lines* show the interfacial polar contacts and salt bridges, and those including the S1/S2 loop arginines are highlighted by ellipses.

TMPRSS2 catalytic residues (H296, D345 and S441) were observed to bind near the S1/S2 cleavage region _681_PRRA**RS**_686_ of SARS-CoV-2 S in 7.5% of the generated models (Extended Data **Fig. 1b**). The corresponding binding affinities were calculated to vary from -14.1 to -11.3 kcal/mol with an average of -12.7 ± 2.0 kcal/mol. **Fig. 2a** displays the most energetically-favorable model where the three arginines in _681_PRRA**RS**_686_ penetrate in the TMPRSS2 catalytic cavity (**Fig. 2a***, right*). Three arginines make salt bridges with acid residues (enclosed in *ellipses*): R682 forms one with TMPRSS2 residue D435, R683 inserts into a pocket containing the TMPRSS2 catalytic aspartate D345, and cleavage site arginine R685 with E299, positioning the scissile bond (*spheres*) near TMPRSS2 catalytic residues S441 and H296.

In the case of furin binding, in contrast to TMPRSS2, 70% of the generated structural models showed the catalytic residues (D153, H194 and S368) stabilized in close proximity of the S1/S2 cleavage region _681_PRRA**RS**_686_ (see Extended Data **Fig. 2**), indicating that binding of furin to the cleavage site was entropically more favorable than that of TMPRSS2. The corresponding binding affinities varied from -16.4 to -11.8 kcal/mol with an average of -14.1±2.3 kcal/mol. The best pose with the catalytic residues facing the S1/S2 site is shown in **Fig. 2b** (and Extended Data **Fig. 2a**), and reveals a number of tight interactions, mostly with S1/S2 loop residues R682 and R683 inserting into negatively charged pockets of furin to enable the cleavage of the SARS-CoV-2 S.

Overall, our analysis shows that either TMPRSS2 or furin can bind the S1/S2 cleavage site and engage in tight intermolecular interactions, in which the basic residues R682 and R683 on the PRRA insert reach out to the catalytic residues of either protease. Binding of either enzyme is accommodated by changes in the local conformations near the cleavage region. However, *in silico* models also suggest that furin binds to the S protein S1/S2 site with higher potency and probability, favored both enthalpically and entropically, compared to TMPRSS2.

### *In silico* screening identifies 4A8 as a potential antibody blocking the S1/S2 cleavage site

SARS-CoV-2 S glycoprotein is the major target of neutralizing mAbs^24,26,27,31-34^. As listed in **Table 1**, cryo-EM studies of antibody-, nanobody- or Fab-bound SARS-CoV-2 S, have been carried out upon substituting the polybasic segment _682_RRAR_685_ by GSAS or SGAG ^22-28,40^ for practical purposes. These mutations would not affect the binding of the Abs that target the RBD (or NTD); but would affect the binding, if any, to the S1/S2 site, as the mutant S lacks the unique character of the highly reactive polybasic segment _682_RRAR_685_. We hypothesized that *in silico* analyses conducted for the wt S protein might capture features that could have been overlooked in the studies carried out with mutants. Therefore, we generated structural models for the intact S as well as its GSAS-mutant using SWISS-MODEL^20^ based on the cryo-EM structures of the dominant conformational states (PDB: 6VSB^15^ and 6VXX^14^); and we tested *in silico* the binding properties of the mAbs listed in **Table 1** to both structures using ClusPro^38^, to examine the possible binding of the Abs near the S1/S2 cleavage site.

Simulations conducted for the GSAS-substituted mutants showed that, not surprisingly, none of the mAbs listed in **Table 1** bound with high affinity to the S1/S2 region, consistent with cryo-EM studies. Multiple binding sites were observed within the RBD and NTD of S1, as well as subunit S2, pointing to possible alternative binding regions, but the vicinity of the mutation site did not attract the mAbs. In contrast, simulations repeated with the wt S protein showed that _682_RRAR_685_ would bind with high affinity the SARS-CoV-2-neutralizing mAb 4A8, and potentially block the S1/S2 cleavage site. 4A8 was identified by cryo-EM to bind to the NTD of the S protein^26^, near residues H145-S150 and G245-R256 (**Fig. 3a**). Notably, our docking simulations also showed that the S epitope and 4A8 paratope observed in cryo-EM indeed engaged in tight interactions (**Fig. 3b**). However, the SAg-like motif E661-R685 also emerged as another equally favorable site in our simulations (**Fig. 3c**). Remarkably, a close overlap between the site occupied by 4A8 and the (otherwise) binding pose of TMPRSS2 or furin was noted, as illustrated for TMPRSS2 in **Fig. 3d**, suggesting that 4A8 might compete with the proteases for binding this site.

**Figure 3:**
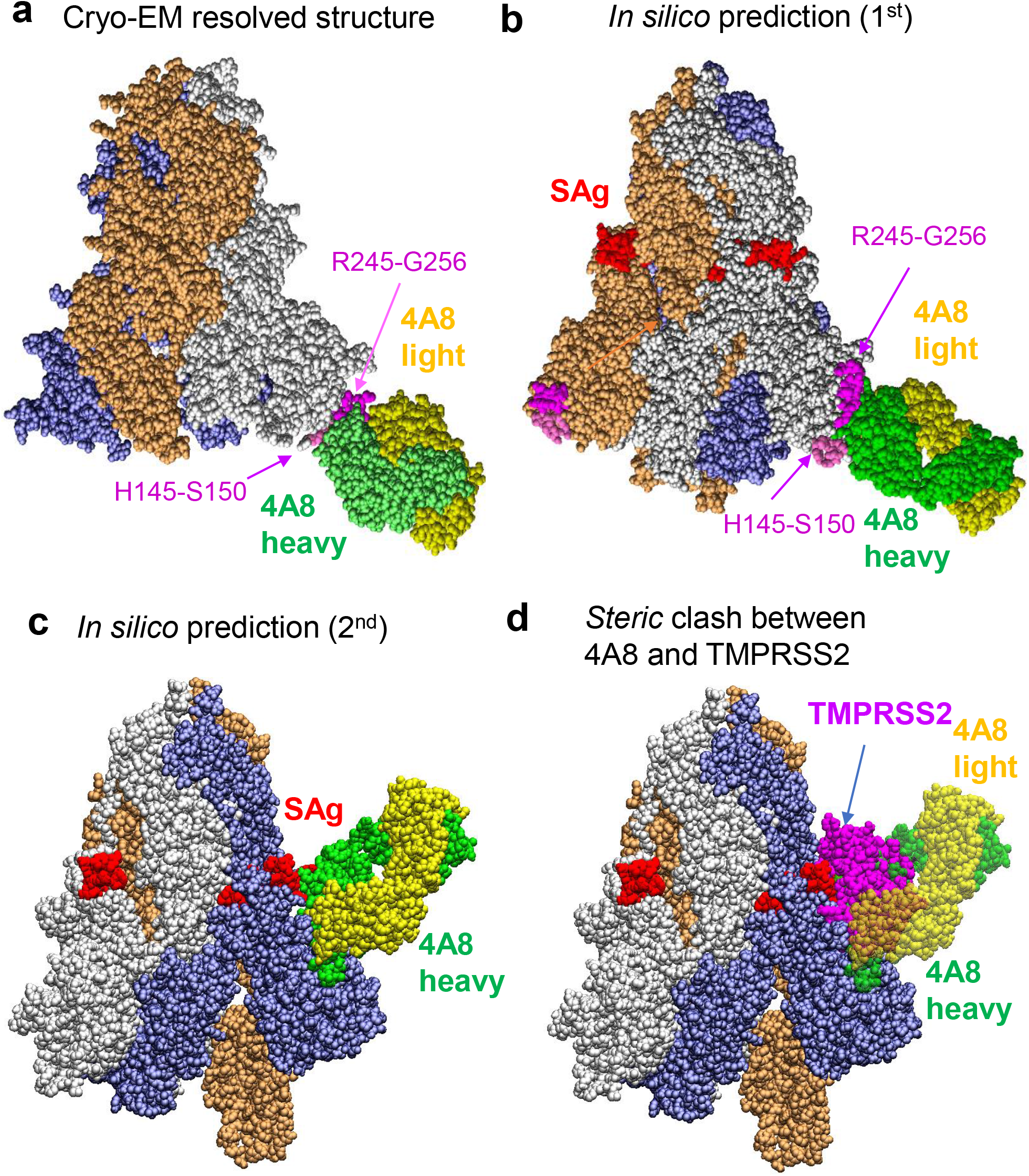
Examination of binding characteristics of SARS-CoV-2-neutralizing mAbs 4A8. **(a)** Cryo-EM structure (PDB: 7C2L)^26^; (**b-c**) Energetically most favorable conformers predicted for the S protein-4A8 complex. The former resembles the cryo-EM structure, involving the same segments, H145-S150 and R245-G256, at the binding epitopes of S. In the latter case the viral SAg-like region which also overlaps with the S1/S2 cleavage site serves as 4A8-binding epitope. (**d)** Competition between 4A8 and TMRPSS2 for binding to the S1/S2 cleavage site, based on the overlap between the binding poses of the two substrates. The diagram is generated by superposing the S-protein of the two complexes predicted *in silico*. The Spike-4A8 complex is generated *in silico* using the SARS-CoV-2 S structure with one RBD in the up conformer (PDB: 6VSB).

To better assess the ability of 4A8 to possibly compete with host cell proteases to bind the S1/S2 site, we evaluated the binding affinity of 4A8 to the S protein in comparison to those of TMPRSS2 and furin. The calculated affinity of 4A8 to bind the S1/S2 site varied from -15.8 to -11.0 kcal/mol with an average of -13.4±2.4 kcal/mol; and its affinity to the NTD site identified by cryo-EM (PDB: 7C2L) varied from -13.8 to -11.2 kcal/mol with an average of -12.5±1.3 kcal/mol. Notably, the affinity of 4A8 to bind the S1/S2 cleavage site is comparable to that of TMPRSS2, and slightly weaker than that of furin. Interestingly, the equilibrium dissociation constants (*K*_d_) measured by biolayer interferometry for the complexes that 4A8 formed with S and with subunit S1 were less than 2.14 nM, and 92.7 nM, respectively^26^. The binding energies (ΔG) based on these respective *K*_d_ values would be -12.3 and -9.6 kcal/mol at 298 K. The computed binding energies show therefore good agreement with the measured *K_d_* for the S protein-4A8 complex. It is interesting to note that binding to S1 was less favorable than that to S protein^26^. This could be attributed to an increased disorder at the “PRRAR” region in the absence of the scaffold provided by the intact S protein, as well as the lack of the contribution of adjoining S2 residues (e.g. S686) to intermolecular association.

Previous studies demonstrated that the mAb 4A8 has strong neutralizing capacities against both authentic and pseudotyped SARS-CoV-2^26^. While we do not exclude the existence of neutralizing effects due to its binding to the NTD site observed in cryo-EM (PDB:7C2L)^26^, our findings suggest that interfering with the access of proteases to the cleavage site S1/S2 could also contribute to its neutralizing effects. Furthermore, the comparable binding affinities of TMPRSS2 and 4A8 mAb competing for the S1/S2 site suggest that mAb blocking of the cleavage site is possible with a sufficiently high-affinity mAb. We next proceed to the identification of more specific, high affinity mAbs that bind the _681_PRRARS_686_ motif.

### Anti-SEB antibody 6D3 is distinguished by its high affinity to bind SARS-CoV-2 S SAg-like region

Because the S residues E661-R685 that enclose the polybasic segment _681_PRRAR_685_ are sequentially and structurally similar to the segment T150-D161 of SEB^8^, we examined if mAbs specific to SEB^35^, could be used for neutralizing SARS-CoV-2 S protein as well. The close proximity of the viral SAg-like region to the S1/S2 proteolytic cleavage site (_685_R-S_686_ peptide bond), suggests that an anti-SEB mAb that cross-reacts with SARS-CoV-2 would have the added potential to block the cleavage site essential to viral entry, apart from its potential modulation of SAg-mediated hyperinflammatory cytokine storm^41^.

Three SEB-associated mAbs, 14G8, 6D3, and 20B1, have been generated as effective blockers of the SAg activity of SEB in an animal model of TSS^35^. Examination of the corresponding crystal structures shows that these antibodies bind different sites on SEB^35^, as illustrated in **Fig. 4a.** Notably, only 6D3 targets the SEB polybasic segment T150-D161 that is sequentially and structurally similar to the SARS-CoV-2 S SAg-like motif. A closeup view shows the tight interaction between the acidic residues E50, D52 and D55 of mAb 6D3 heavy chain and four basic residues of SEB (**Fig. 4b**).

**Figure 4:**
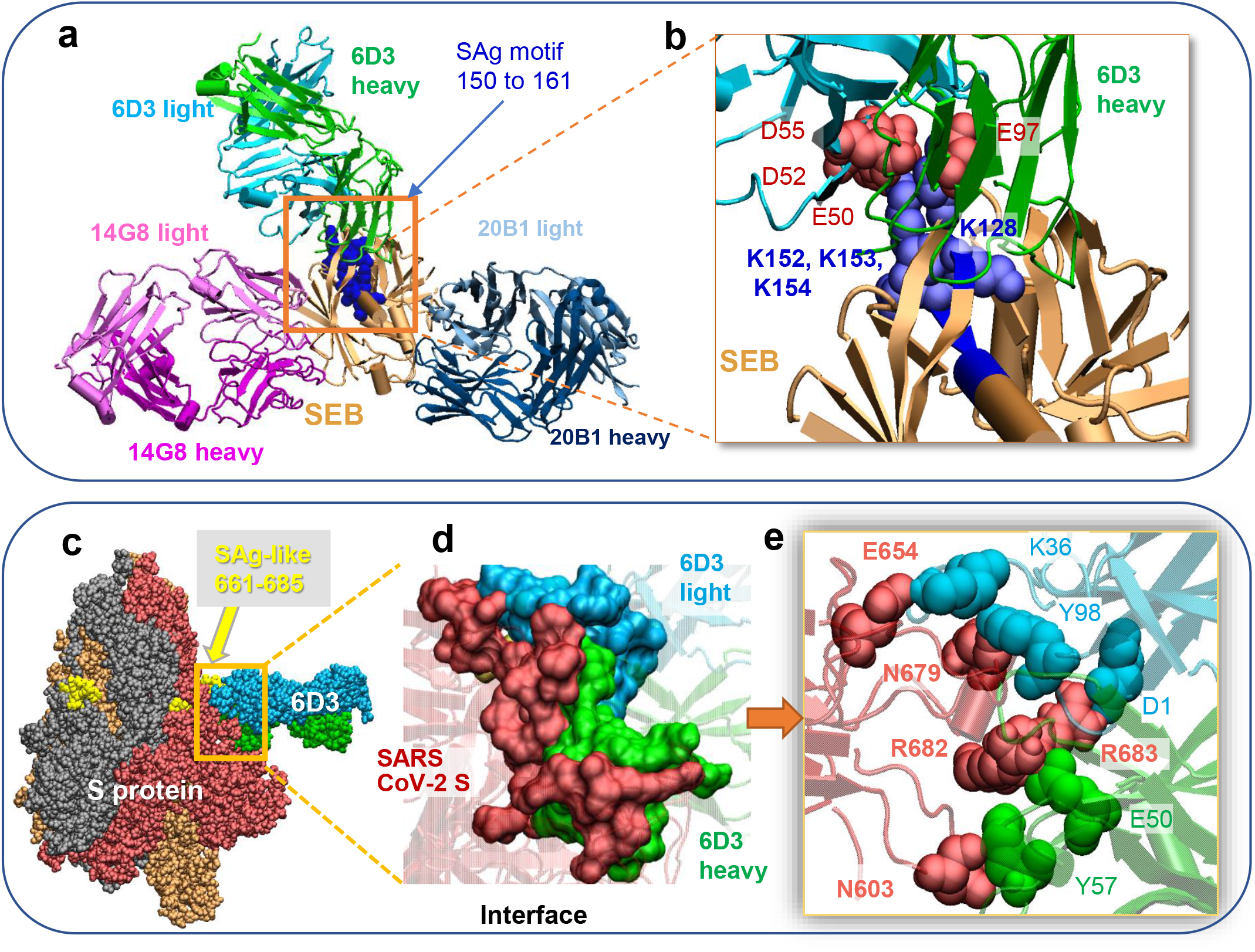
SEB-associated mAb 6D3 exhibits a high affinity to bind to the furin-cleavage site of SARS-CoV-2 S protein, potentially interfering with the S1/S2 cleavage by furin or TMPRSS2. (**a**) Binding pose of three SEB-neutralizing Abs (mAbs 6D3, 14G8, and 20B1) onto SEB. The diagram is generated by superposing the crystal structures (PDB IDs 4RGN and 4RGM) resolved for the complexes^35^. SEB is colored beige, with its SAg motif _150_TNKKKATVQELD_161_ highlighted in *blue* s*pace-filling*. (**b**) Interface between 6D3 and SEB SAg motif. Heavy and light chains of 6D3 are colored *green* and *cyan*, respectively. Overall (**c**) and close-up (**d**) views of the complex formed between the S protein and anti-SEB mAb 6D3 and corresponding interfacial interactions engaging the arginines in the PRRA insert. SARS-CoV-2 S interfacial residues include I210-Q218, N603-Q607, E654-Y660 and A688-I693, and the SAg motif residues Y674, T678-R683. 6D3 interfacial residues include A24-K33, E50, D52, S54, D55, Y57, N59, K74-T77, and A100-A104 in the heavy chain, and D1, I2, Q27, N31-F38, Y55, W56, and D97-Y100 in the light chain. The Spike-4A8 complex is generated *in silico* using SARS-CoV-2 S structure with one RBD in the up conformer (PDB: 6VSB).

As expected, docking simulations with S protein and these anti-SEB mAbs also showed that only 6D3 was able to bind to the SARS-CoV-2 S SAg motif (**Fig. 4c-e**), consistent with 6D3 binding to the precise SEB fragment that aligns with the spike SAg-like motif. Our computational analysis further predicted the 6D3 Ab to have a greater affinity for binding this polybasic region compared to TMPRSS2, and an affinity comparable to that of furin. The 6D3 binding affinity was -14.2 ± 2.3 kcal/mol based on a cluster of conformers that exhibited similar binding features (see Supplemental Methods). Notably, acidic residues E50, D52 and D55 from the heavy chain of 6D3 were again found to interact with polybasic insert PRRA in SARS-CoV-2 S, with R682 and R683 playing a central role as will be elaborated below; but interfacial contacts were quite distributed, involving other SARS-CoV-2 S amino acids such as E654, N603 and N679 interacting with either the heavy or light chains of 6D3 (**Fig. 4e**). Among those 6D3-interacting S residues, N603 has been identified as an N-linked glycan site by site-specific glycan analysis of SARS-CoV-2 S^42^. To further investigate if glycan sequons may interfere with Ab 6D3 binding, we aligned our spike-6D3 complex model against the glycosylated spike^43^; no steric overlap was observed between Ab 6D3 and glycan sequons. The N603-linked glycan might even be involved in the recognition by 6D3 rather than obstructing its binding.

Based on these results, we proposed that 6D3 may have the capacity to attenuate, if not completely block, SARS-CoV-2 infection upon effectively competing with the host cell proteases TMPRSS2 and furin to bind to the cleavage site, and thus prevent the proteolytic cleavage of the S protein monomers that is essential to S protein priming for viral entry.

### Anti-SEB antibody, 6D3, inhibits SARS-CoV-2 infection

Our *in silico* analyses led to the identification of two mAbs (4A8 and 6D3) that may potentially block SARS-CoV-2 S1/S2 cleavage site. 4A8 has been demonstrated in previous work to possess strong neutralizing capacities against both authentic and pseudotyped SARS-CoV-2^26^. We investigated here whether the anti-SEB antibody 6D3 also had a similar capacity. To this end, we tested the ability of 6D3 to inhibit SARS-CoV-2 infection in an *in vitro* cell culture infection system. Antibodies were incubated with SARS-CoV-2 for 1 hour and then added to plated Vero-E6 cells. At 48 hours post infection, we analyzed viral infection by immunofluorescence using antibodies against dsRNA or SARS-CoV-2 S protei. (**Fig. 5** and Extended Data **Fig. 3**). We found that 6D3 significantly inhibited viral infection, as measured by the percentage of dsRNA positive cells, at concentrations of 0.8, 4 and 20 μg/ml of antibody (**Fig. 5a-b** and Extended Data **Fig. 3a**). Furthermore, in an independent set of experiments, we found that 6D3 significantly inhibited viral infection, as measured by the percentage of spike positive cells, at concentrations of 4, 20 and 40 μg/ml of antibody, while there was a trend for inhibition at 0.16 and 0.8 μg/ml of antibody (**Fig. 5c-d** and Extended Data **Fig. 3b**).

**Figure 5:**
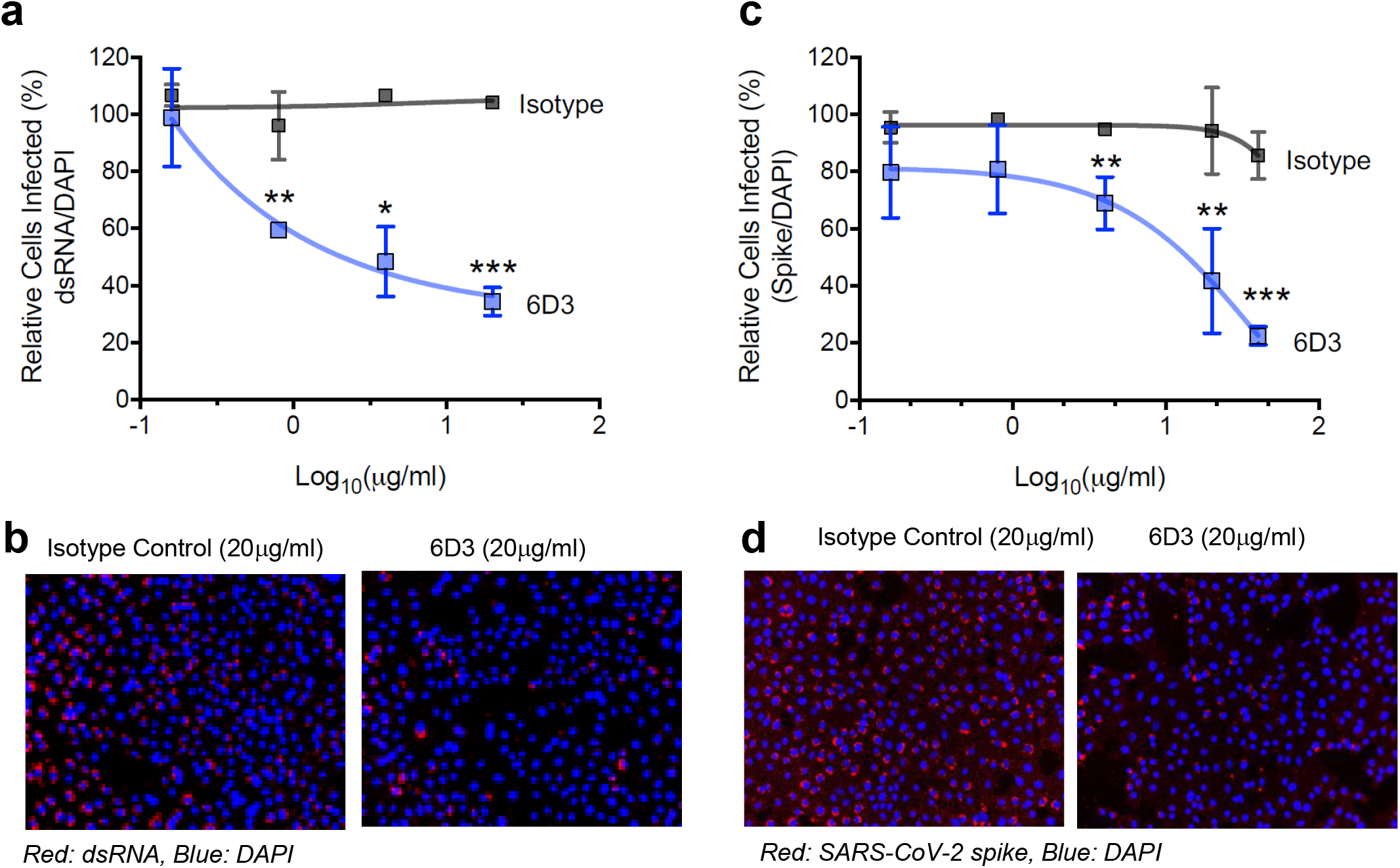
Monoclonal antibody 6D3 prevents SARS-CoV-2 infection. 6D3 or isotype control antibodies (at indicated concentrations) were incubated with virus (100 PFU/well) for 1 hour at room temperature before addition to Vero-E6 cells (5×10^3^ cells/well). 48 hours post infection cells were fixed and stained for dsRNA or SARS-CoV-2 spike protein. (**a**) Quantification of the percentage of infected cells per well by dsRNA staining. Data are representative of two independent experiments. (**b**) Representative fluorescence images of 6D3-mediated inhibition of virus infection (dsRNA). (**c**) Quantification of the percentage of infected cells per well by spike staining. (**d**) Representative fluorescence images of 6D3-mediated inhibition of virus infection (spike). Data were analyzed by t test (6D3 vs. isotype control) with multiple testing correction (FDR). See also Extended Data **Fig. 3** for detailed results as a function of 6D3 concentration.

Combined with *in silico* modeling, these results indicate that 6D3 can block the protease cleavage site of SARS-CoV-2 Spike to prevent viral entry in a concentration dependent manner, and confirm our previous finding that SARS-CoV-2 Spike possesses a SAg-like structure similar to SEB. 6D3 may therefore have a dual potential to block SARS-CoV-2 infection as well as SAg-mediated T cell activation and hyperinflammation, with implications for treating cytokine storm during severe COVID-19 and MIS-C/A diseases.

### An acidic residue cluster at VH CDR2 is the hallmark of Abs targeting the furin-like cleavage site

Our study pointed to the distinctive ability of two mAbs, 6D3 and 4A8, to bind the S1/S2 cleavage site, while other mAbs (in **Table 1**) did not show such a binding propensity. We investigated which sequence/structure features distinguish these two mAbs from all others. Abs target viruses mainly through their three complementarity determining regions (CDR1-3) in the variable domains, especially those in the heavy chains^44^. **Fig. 6a** compares the sequences of the variable heavy (VH) chains of SARS-CoV-2 S-associated mAbs, as well as those of the three mAbs associated with SEB. CDR3s exhibit the largest sequence variation, in accord with their role in conferring specificity. However, comparison of the sequences reveals a unique feature that distinguishes 6D3 and 4A8, mainly a poly-acidic cluster at their CDR2. Specifically, the 6D3 CDR2 possesses three acidic residues E50, D52, and D55, already noted above to enable binding to not only the SAg-like motif of SEB and SARS-CoV-2, but also the precise cleavage-site on the S protein. Likewise, mAb 4A8 has four acidic residues D52, E54, D55 and D57, compared to none in the majority of the RBD-targeting mAbs (**Fig. 6a**).

**Figure 6:**
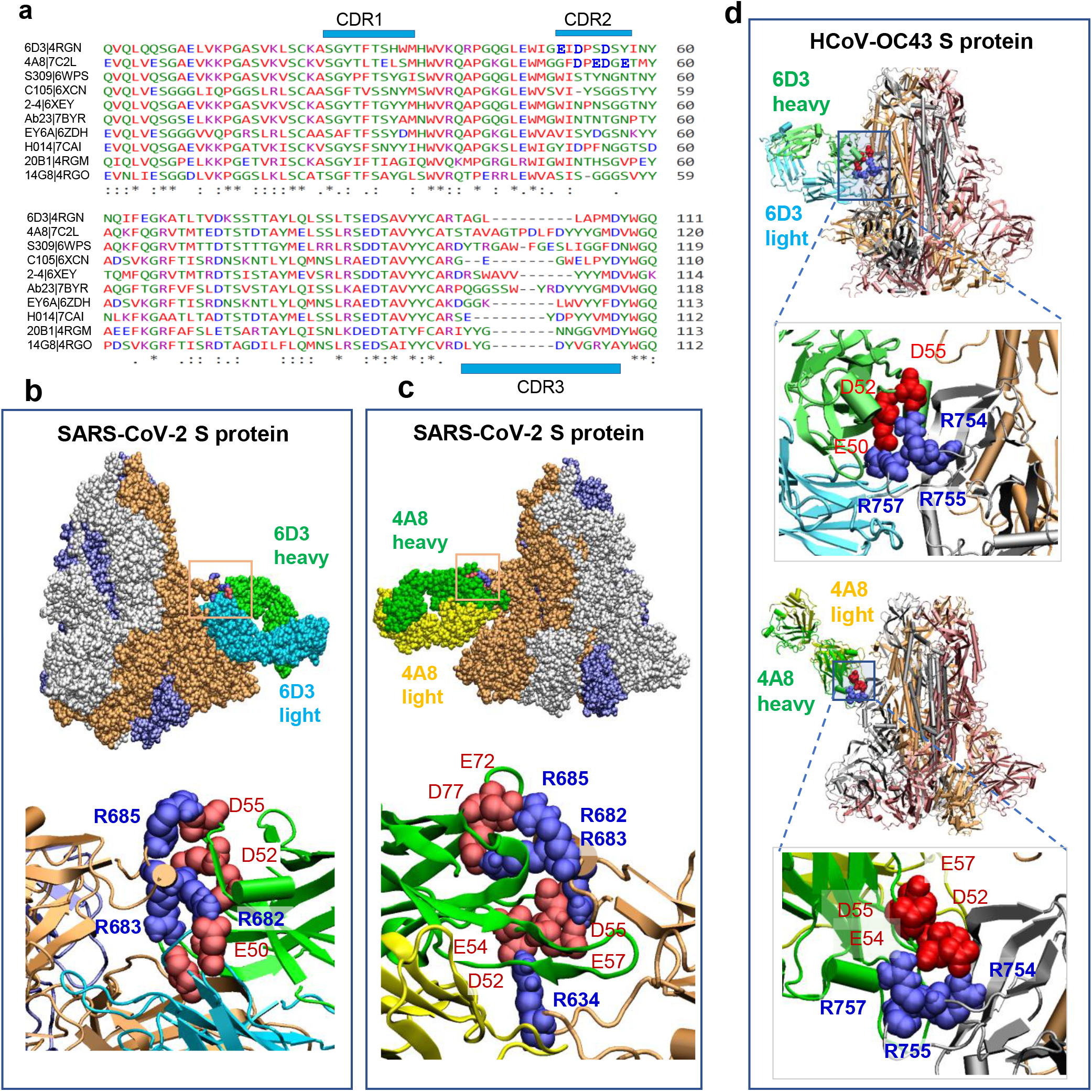
Polyacidic residues in the CDR2 of the mAbs 6D3 and 4A8 heavy chains play a major role in blocking the furin-like cleavage site of HCoV S protein. (**a**) Multiple sequence alignment of the VH domain of anti-SEB Abs (6D3, 14G8 and 20B1) and anti-SARS-CoV-2 S Abs (listed in the *left column*). Using Ab 4A8 as the reference, the residue ranges of the three CDRs are: CDR1 (residues 25 to 32), CDR2 (51 to 58), and CDR3 (100 to 116)^26^, as indicated by the *blue bars*. (**b**) Overall and close-up views of the complex and interfacial interaction of SARS-CoV-2 S protein and 6D3 antibody. Note that three acidic residues from CDR2 interact with the basic residues R682, R683 and R685 of the S protein. (**c**) Same as **b**, repeated for 4A8. A cluster of acidic residues from CDR2 forms a network of interactions with the S protein R682, R683 and R685. The complexes in panels **b** and **c** are generated *in silico* using SARS-CoV-2 S structure with three RBDs in the down conformer (PDB: 6VXX). The residues belonging to the Abs are labelled in *lightface*, those of the S protein in *boldface.* HCoV-OC43 encodes a S1/S2 furin-like cleavage site at _**754**_RRA**R**↑**G**_758_. (**d**) Overall and close-up views of the complex formed, and interfacial interactions, between the HCoV-OC43 S protein (with S1/S2 furin-like cleavage site _**754**_RRA**R**↑**G**_758_) and the Ab 6D3 (*top*) and 4A8 (*bottom*). Note that three acidic residues from CDR2 interact with R754, R755 and R757 in hCoV-OC43 S protein. The residues belonging to the Abs are labelled in *lightface*, those of the HCoV in *boldface.*

A poly-acidic CDR2 at the VH chain thus emerges as a hallmark of the mAbs that target the polybasic furin-like cleavage site. As shown in **Fig. 6b** and **c**, these acidic residues facilitate Ab-spike complexation through salt bridges formed with the basic residues (R682, R683 and R685) in the _680_SPRRARSV_687_ segment, the central component of a typical furin-cleavage site^36^, thus blocking the access of proteases to the S1/S2 cleavage site.

As shown in **Fig. 1b**, the polybasic insertion of SARS-CoV-2 is not shared by other SARS-family βCoVs, but is found in common cold HCoVs HKu1 and OC43, and in MERS. Present findings strongly suggest that mAbs 6D3 and 4A8 may also target these other HCoVs that encode a furin-cleavage site. To test this hypothesis, we generated a structural model for HCoV-OC43 S protein based on the cryo-EM structure resolved for OC43 (PDB: 6NZK)^45^ using SWISS-MODEL^20^. We then investigated the binding properties of the Abs 6D3 and 4A8 to that structure with docking simulations. The most stable (highest binding affinity) poses resulting from our simulations are presented in **Fig. 6d.** Both 6D3 and 4A8 are therefore predicted to bind the S1/S2 furin cleavage site of HCoV-OC43 S. Again, poly-acidic residues in CDR2 play a primary role in binding to the cleavage site of HCoV-OC43 S. These findings underscore the potential effectiveness of Abs that target the S protein S1/S2 cleavage site, and their cross-reactivity between the S proteins of SARS-CoV-2 and other selected HCoVs.

## Discussion

### A new strategy for combatting SARS-CoV-2: repurposing of antibodies that target the S1/S2 cleavage site

There is an urgent need to design effective therapeutics that inhibit SARS-CoV-2 infection and suppress the hyperinflammatory cytokine storm in severe COVID-19 and MIS-C/A patients. The most effective measure against SARS-CoV-2 is vaccination, and multiple SARS-CoV-2 vaccines are about to be released following several successful phase III trials^46^. However, the duration of the immune protection, their ability to protect special populations across all age groups, as well as the capacity to produce sufficient doses of safe and effective vaccines remain to be determined^46^, highlighting further the urgent need to discover effective therapeutics.

SARS-CoV-2 S is the main determinant of cell entry and the major target of neutralizing Abs^24,26,27,31–34^. Binding to receptor ACE2 is an essential first step in establishing infection; therefore the majority of COVID-19 Ab therapies under investigation are designed to target the S protein RBD, while other potential neutralizing epitopes have also been found^23,24,26,27,31–34^. Given the high glycosylation and antigenic variability of SARS-CoV-2 S^47^, a combination of mAbs that target multiple sites and multiple conformations of SARS-CoV-2 S, is likely the most effective strategy. Besides blocking ACE2 binding, distinct neutralizing mechanisms have been proposed, including Ab-dependent cell cytotoxicity and phagocytosis^24^ and restraining the conformational changes of SARS-CoV-2 Spike^26^.

Proteolytic cleavage of SARS-CoV-2 S is the second critical step, succeeding ACE2 binding, in the life cycle of SARS-CoV-2. Host proteases shown to prime the SARS-CoV-2 S include cell surface TMPRSS2, furin, and endosomal cathepsins^16,48^. TMPRSS2 and furin inhibitors have been found to block cell entry of SARS-CoV-2^16,49^. Unlike TMPRSS2, furin is a ubiquitous proprotein convertase and is required for normal development and function^50^. Thus, repurposing of Abs that block the S1/S2 cleavage site is an attractive alternative solution that avoids effects on the (other) biological activities of TMPRSS2 and furin.

It is well known that the SARS-CoV-2 spike is heavily glycosylated, and the possible interference of glycans with Ab binding is a plausible consideration. Notably, 6D3- or 4A8-binding did not give rise to steric clash with the N-linked glycan sequons near the S1/S2 site (e.g. N603, or N657/N658 as reported^42^). In addition SARS-CoV-2 S was predicted to be O-glycosylated at S673, T678 and S686 near the S1/S2 cleavage site^51^, yet to be confirmed by experiments^42,52^. Therefore, mAbs may possibly escape the glycan shields and target directly the S1/S2 site of SARS-CoV-2 S as the host proteases do.

### The ability of the polybasic insert to bind antibodies may have escaped prior cryo-EM studies with mutant S protein

The detailed atomic structure of the S1/S2 cleavage site of SARS-CoV-2 S is yet to be determined. It has been a challenge to resolve this region in cryo-EM studies of HCoV S proteins. First, pre-activation of HCoV S during protein preparation results in a mixture of cleaved and un-cleaved spikes^53^. Second, local conformational changes near the S1/S2 region may differ between cleaved and intact structures, as observed in influenza viruses^54^. Third, multiple conformations, if not a disordered state, may exist near that region, as indicated by microseconds molecular dynamics simulations and *ab initio* modeling^55^. Therefore, it has been difficult to resolve that region, and the majority of cryo-EM studies of SARS-CoV-2 S protein complexed with Abs have resorted to mutant S proteins where the _682_RRAR_685_ segment has been replaced by GSAS or SGAG^22–28,40^ (**Table 1**). These ‘mutant spikes’ may have precluded the discovery of binding of Abs to the S1/S2 site. Molecular modeling and simulations provided insights into the interactions at this region, including those with proteases and other receptors^8,55,56^. We undertook such modeling studies (**Figs 2, 3, 4c-e** and **6**), and performed live-virus experiments (**Fig. 5** and Extended Data **Fig. 3**) to investigate the Abs that possibly interfere with the binding of host cell proteases TMPRSS2 and furin. To our knowledge, we identified the first two antibodies (6D3 and 4A8) that may block the S1/S2 cleavage site in SARS-CoV-2 S protein.

### 6D3 is a repurposable anti-SEB mAb that targets the S1/S2 site and inhibits viral infection

6D3 is an antibody originally discovered for neutralizing the superantigenic bacterial toxin SEB. Here we are proposing its use as repurposable antibody against SARS-CoV-2 S protein, by virtue of its ability to bind a sequence motif shared between SEB and S protein. Our recent study indeed revealed the high sequential and structural similarity between SARS-CoV-2 S amino acids E661-R685 and SEB amino acids T150-D161, which may contribute to hyperinflammation and MIS-C/A pathogenesis through a SAg-induced immune activation^8^. This view was supported by the observations that the clinical and laboratory features observed in MIS-C and severe COVID-19 patients were similar to those of toxic shock syndrome (TSS) caused by bacterial toxins such as SEB^9^; and adult patients with severe Covid-19 displayed TCR skewing typical of SAg-induced immune response^8^. Therefore, we hypothesized that mAbs that target the homologous segment of SEB, if any, could be repurposed as SARS-CoV-2 Abs. Among the three mAbs discovered against SEB^35^, 6D3 was the only one specific to the region of interest (**Fig. 4a-b**), and computations and experiments corroborated our hypothesis that this anti-SEB mAb could bind the SARS-CoV-2 S protein.

It remains to be seen if 6D3 can effectively interfere with the superantigenic activity of the S protein and possibly suppress the hyperinflammation and cytokine storm associated with MIS-C/A. However, one other feature that caught our attention was the fact that this SAg-like segment (that binds 6D3) also overlapped with the S1/S2 cleavage site, a furin-like cleavage site characteristic of SARS-CoV-2 (and MERS and HCovs HKU1 and OC43; see **Fig. 1b**). Furin-cleavage sites usually involve ~20 residues, with eight residues in the vicinity of the cleavage site playing a central role^36^. In the case of SARS-CoV-2, the segment _680_SPRRAR↑SV_687_ forms this central component of the furin cleavage site at the S protein. Simulations indeed showed strong interactions (salt bridges) formed between 6D3 VH CDR2 (distinguished by a stretch of acidic residues) and the polybasic segment _682_RRAR_685_ of the S protein (**Figs 4c-e** and **6b**), and *in vitro* assays confirmed that 6D3 inhibited viral entry (**Fig. 5** and Extended Data **Fig. 3**).

Notably, 4A8, previously shown to inhibit SARS-CoV-2 infection^26^, also possesses the same polyacidic properties at its VH CDR2 (**Fig. 6a** and **c**), which distinguishes 6D3 and 4A8 from all other SARS-CoV-2-S-associated mAbs. These results suggest that the effectiveness of 4A8 may have originated from its high affinity to bind the polybasic insert at the furin-like cleavage site (alongside possible binding to NTD observed in cryo-EM^26^). Structural characterization of the wt S protein complexed with 4A8 could shed light onto this possibility.

### mAbs containing a cluster of acidic residues at their VH CDR2 may mitigate viral infections caused by CoVs that contain furin-like cleavage sites

HCoVs include three highly pathogenic viruses, SARS-CoV-2, SARS-CoV and MERS, and four circulating endemic viruses (HCoV-NL63, HCoV-229E, HCoV-OC43 and HKU1) which cause mild to moderate upper respiratory diseases^10-12^. Interestingly, many individuals who have not been exposed to SARS-CoV-2 possess SARS-CoV-2 Spike reactive T cells, due to cross-reaction of immune responses generated against other HCoV strains^57,58^. Cross-reactive antibodies between human βCoV strains have also been identified, including those between SARS and SARS-CoV-2^59,60^. Indeed, SARS monoclonal antibody S309 can potently neutralize both SARS and SARS-CoV-2^24^. Furthermore, the effectiveness of IVIG^4-6^, may, in part, be due to the presence of cross-reactive antibodies against other HCoV stains. These findings raise the exciting possibility of designing wide spectrum Abs with cross-reactivity among HCoVs. The two Abs (6D3 and 4A8) identified in this study to bind the PRRAR insert and cleavage site may potentially act as wide spectrum Abs to block S1/S2 cleavage site in HCoVs that encode furin-like cleavage sites (**Fig. 6**), providing benefit beyond the current pandemic. Previous work also showed the role of CDR2-binding in neutralizing a toxin^61^. The hallmark poly-acidic residues in the CDR2 of VH may be exploited as a benchmark to sort out mAbs able to target the SARS-CoV-2 furin cleavage site.

### Alternative strategies targeting the S1/S2 site in the light of these repurposable mAbs

Based on the scaffold of 6D3 or 4A8 heavy chain, mini-proteins may be designed to target SARS-CoV-2, MERS, HCoV-OC43 or HKU1, to block CoV entry. Notably, designed *de novo* mini-proteins have been shown to block ACE2 binding, based on the scaffold of ACE2^62^. Very recently, neuropilin-1 (NRP1) has been identified as a host factor for SARS-CoV-2 infection, bound to the _681_RRAR_685_ segment^30^. Remarkably, blockade of this interaction by RNAi or mAb against NRP1 significantly reduced *in vitro* SARS-CoV-2 cellular entry^29,30^. We anticipate that 6D3 or 4A8 can effectively block the binding of NRP1. At present, no clinical treatments or prevention strategies are available for HCoVs^12^. Our work may lead to an improved understanding of coronavirus immunity, facilitating future studies to understand mechanisms of antibody recognition and neutralization, and help screen SARS-CoV-2 Abs for treatment of COVID-19. These findings also raise exciting possibilities of designing novel therapeutic approaches using a combination treatment of Abs 6D3 or 4A8 to treat severe COVID-19 and MIS-C/A patients. Likewise, cocktail treatment including furin-like cleavage site-targeting Abs with RBD-targeting neutralization Abs may provide complementary protection against SARS-CoV-2 pathogenesis.

## Acknowledgements

We gratefully acknowledge support from the NIH awards 3RO1AI072726-10S1 (to MA) and P41GM103712 (to IB) and a MolSSI COVID-19 Seed Software Fellowship (to JK). This study was in part funded with resources provided by US Veterans Affairs Merit Review Award 5I01 BX003741 (to BCF). BCF is an attending at the U.S. Department of Veterans Affairs - Northport VA Medical Center, Northport, NY. The contents of this study do not represent the views of VA or the United States Government.

## Author Contributions

MHC, IB, and MA designed and supervised the work. MHC did the computations, assisted by JK. IB and MHC analyzed the computational results. VA and GG performed the *in vitro* viral infections. ABO and RAP analyzed the experimental *in vitro* data. BCF interpreted experimental data. MHC, RAP, MNR, MA and IB wrote the manuscript.

