## Supplemental methods and figures for "A monoclonal antibody against staphylococcal enterotoxin B superantigen inhibits SARS-CoV-2 entry *in vitro*"

**Author Footnotes**

[*] These senior authors contributed equally

### Supplemental Methods

**Structural data for SARS-CoV-2, human TMPRSS2 and furin**. SARS-CoV-2 (residues A27-D1146; UniProt ID: P0DTC2) spike models were generated using SWISS-MODEL[1], based on the resolved SARS-CoV-2 Spike glycoprotein structures of SARS-CoV-2 in different conformational states (PDBs: 6VSB[2] and 6VXX[3]). The missing loops in the crystal structures, were built using the well-established libraries of backbone fragments[4] and constraint space *de novo* reconstruction of the backbone segments[5]. The catalytic domain of human TMPRSS2 (residues N146-D491; UniProt ID: O15393) was constructed using SWISS-MODEL[1], based on the crystal structure of serine protease hepsin (PDB: 5CE1). A crystal structure of human furin (Y110-A408; P09958) was used as is (PDB: 5JMO)[6].

**Generation and assessment of SARS-CoV-2 Spike and protease complex models.** To investigate priming of the S1/S2 site of SARS-CoV-2 Spike, we performed protein-protein docking analysis of TMPRSS2 or furin with SARS-CoV-2 Spike in the pre-fusion state. Using docking software ClusPro[7], we constructed *in silico* a series of SARS-CoV-2 Spike and protease complexes. SARS-CoV-2 Spike was set as receptor and protease as ligand. Residues in the proximity of the cleavage site from SARS-CoV-2 Spike (T676 to V687) were set as attractor sites of receptor, and the catalytic residues from TMPRSS2 (H296, D345 and S441) or furin (D153, H194 and S368) were set as attractor sites for ligand. For each complex, we obtained 30 clusters of conformations, upon clustering ~800 models generated by ClusPro. The clusters were rank-ordered by cluster size[7] as recommended, and representative members from top-ranking clusters were further examined and refined Mainly, protein-protein binding free energies were calculated using PRODIGY[8]; and mutagenesis and sculpting wizards in PyMOL 2.3.0 (Open Source version)[9] were used to interactively refine rotamers and interactions, respectively.

**Monoclonal antibodies binding to SARS-CoV-2 Spike.** SEB-associated monoclonal antibodies 14G8, 6D3 and 20B1 were taken from the crystal structures of SEB bound to two neutralizing Abs, 14G8 and 6D3 (PDB: 4RGN), and one neutralizing Ab, 20B1 (PDB: 4RGM). SARS-CoV-2 S-associated neutralizing Abs were taken from the crystal structures listed in **Table 1**. Ab-binding poses were predicted using ClusPro[7]. Selective mAb-Spike complexes were further refined by molecular dynamics simulations performed by HADDOCK 2. 2[10]. Binding free energies were calculated using PRODIGY[8]. Multiple sequence alignment of the variable heavy chain domain of anti-SEB Abs (6D3, 14G8 and 20B1) and anti-SARS-CoV-2 S Abs were generated by Clustal Omega[11].

***In vitro* viral inhibition assays.** SARS-CoV-2 viral assays were performed in UCLA BSL3 high containment facility, following previous procedure[12]. Vero-E6 [VERO C1008 (ATCC@ CRL-1586™)] cells were obtained from ATCC and cultured at 37°C with 5% $CO_2$ in EMEM growth media with 10% fetal bovine serum and 100 units/ml penicillin. SARS-CoV-2 Isolate USA-WA1/2020 was obtained from BEI Resources of National Institute of Allergy and Infectious Diseases (NIAID). Mouse Fab 6D3 (IgG2b) was generated as[13]. Vero-E6 cells were plated in 96-well plates ($5x10^3$ cells/well). 6D3 IgG2b or mouse IgG2b isotype control (Bio X Cell) were incubated with virus (100 PFU/well) for 1 hour at room temperature prior to addition to cells. After 48 hours post-infection the cells were fixed with methanol for 30-60 minutes in -20 °C. Cells were washed 3 times with PBS and permeabilized using blocking buffer (0.3% Triton X-100, 2% BSA, 5% Goat Serum, 5% Donkey Serum in 1 X PBS) for 1 hour at room temperature. Subsequently, cells were incubated with mouse anti-dsRNA antibody (Absolute Antibody, 1:200) or anti-SARS-CoV-2 spike antibody (Sino Biological, 1:200) at 4°C overnight. Cells were then washed 3 times with PBS and incubated with fluorescence conjugated secondary antibody: Goat anti-mouse IgG Secondary Antibody, Alexa Fluor 555 (Fisher Scientific, 1:1000) for 1 hour at room temperature. Nuclei were stained with DAPI (4',6-Diamidino-2-Phenylindole, Dihydrochloride) (Life Technologies) at a dilution of 1:5000 in PBS for 10 minutes. Cells were analyzed by fluorescence microscopy. Five images per well were quantified for each condition. Data were analyzed by T test (6D3 vs. isotype control) with multiple testing correction (Benjamini, Krieger and Yekutieli FDR test).

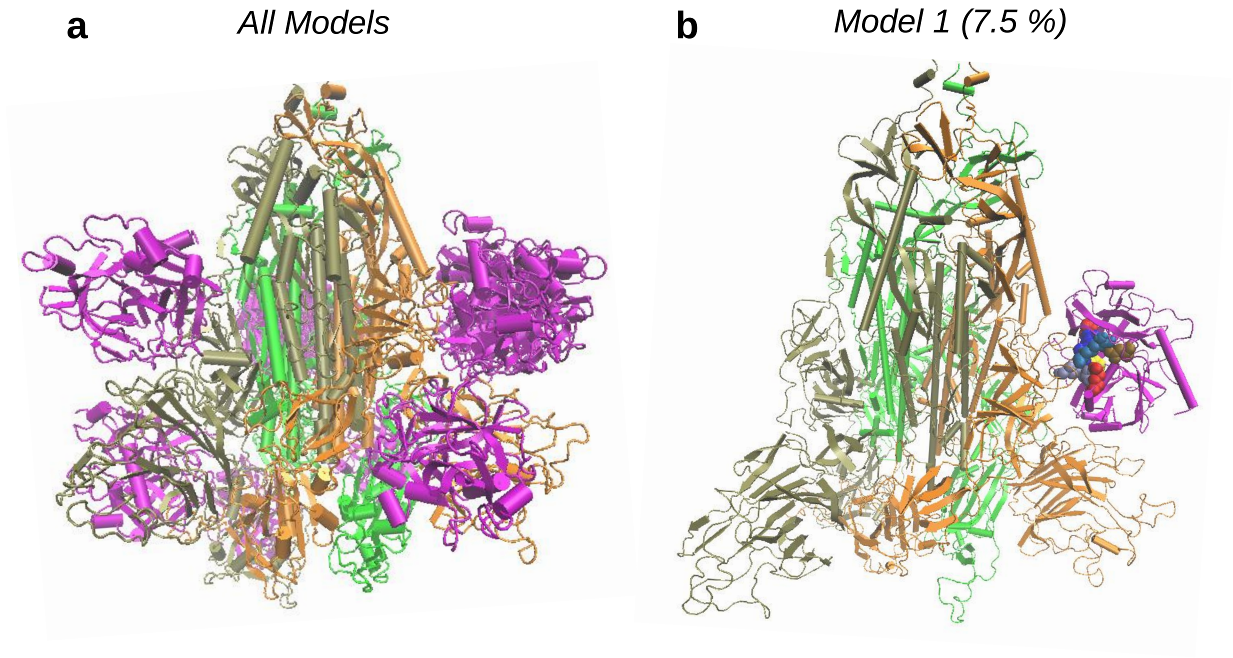


**Extended Data Fig. 1**: **Binding of TMPRSS2 to the SARS-CoV-2 Spike (S) protein yields an ensemble of conformers including a structural model where the protease binds to the PRRA insert**. (**a**) Overlay of models generated by the protein docking software ClusPro [7]. The three subunits of the S protein are colored *tan*, *orange* and *green*; alternative poses of TMPRSS2 are shown in *magenta*. (**b**) Model 1 where TMPRSS2 catalytic residues are positioned in close proximity of the S1/S2 cleavage site. Three basic residues, R682, R683, and R685 from the S protein, are shown as van der Waals (vdW) balls in different shades of *blue*; the acidic residues of TMPRSS2 which form salt bridges with these three basic residues are displayed in *red* vdW balls with catalytic residue D345 in a *darker red* and catalytic serine residue S441 is shown as *yellow* vDW balls.

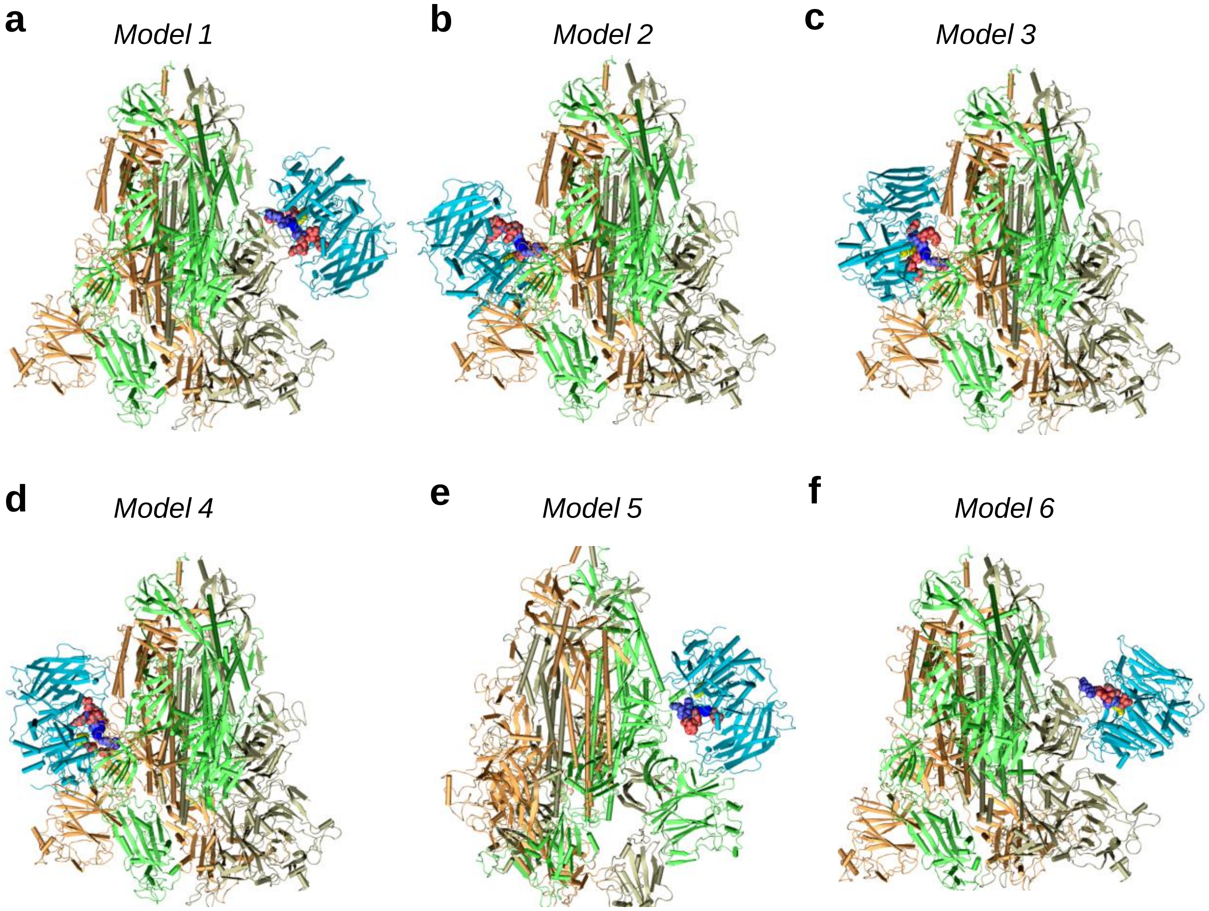


**Extended Data Fig. 2: Structural models generated for SARS-Cov-2 S protein complexed with furin.** Six models, labeled *Model 1* to *Model 6* representative of clusters formed by top-ranking conformers are displayed. In all models, the catalytic residues (D153, H194 and S368) of furin are in close proximity to the cleavage site $_{685}$RS$_{686}$ of spike. The subunits from the S protein are colored *tan*, *orange* and *green*; and furin is in *cyan* cartons. Three basic residues R682, R683, and R685 from spike are shown in *blue* vdW balls; the acidic residues which form salt bridges with these three basic residues from spike are displayed in *red* vdW balls. Note that furin has multiple acidic residues that form intermolecular salt bridges in multiple poses: D153, D154, E236, D258, D264, D306, and E331, and the close proximity of the S1/S2 site is highly favorable, both energetically and entropically. *Model 5*, found to be most favorable energetically, is shown in **Fig. 2b**.

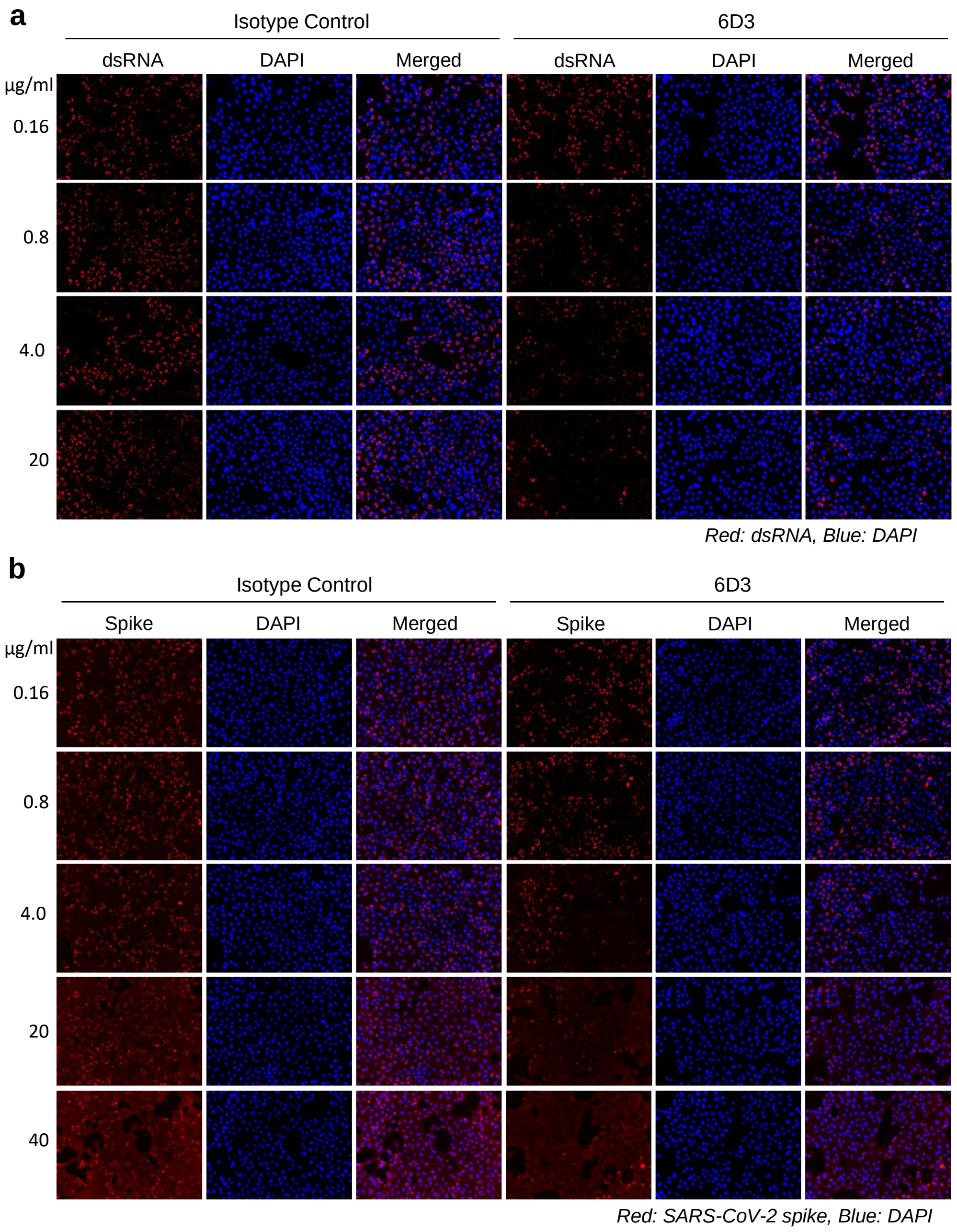


**Extended Data Fig. 3: Monoclonal antibody 6D3 prevents SARS-CoV-2 infection.** 6D3 or isotype control antibodies (at indicated concentrations) were incubated with virus (100 PFU/well) for 1 hour at room temperature before addition to Vero-E6 cells ($5x10^3$ cells/well). 48 hours post infection cells were fixed and stained for dsRNA or SARS-CoV-2 spike protein. **(A)** Representative fluorescence images of 6D3 mediated inhibition of virus infection as measured by dsRNA staining. **(B)** Representative fluorescence images of 6D3 mediated inhibition of virus infection as measured by SARS-CoV-2 spike staining.
